# Structural Adaptation of Darunavir Analogs Against Primary Resistance Mutations in HIV-1 Protease

**DOI:** 10.1101/406637

**Authors:** Gordon J. Lockbaum, Florian Leidner, Linah N. Rusere, Mina Henes, Klajdi Kosovrasti, Ellen A. Nalivaika, Akbar Ali, Nese Kurt Yilmaz, Celia A. Schiffer

**Author notes:** **Corresponding Author** Celia A. Schiffer, Nese Kurt Yilmaz.

## Abstract

HIV-1 protease is one of the prime targets of agents used in antiretroviral therapy against HIV. However, under selective pressure of protease inhibitors, primary mutations at the active site weaken inhibitor binding to confer resistance. Darunavir (DRV) is the most potent HIV-1 protease inhibitor in clinic; resistance is limited, as DRV fits well within the substrate envelope. Nevertheless, resistance is observed due to hydrophobic changes at residues including I50, V82 and I84 that line the S1/S1’ pocket within the active site. Through enzyme inhibition assays and a series of 12 crystal structures, we interrogated susceptibility of DRV and two potent analogs to primary S1’ mutations. The analogs had modifications at the hydrophobic P1’ moiety to better occupy the unexploited space in the S1’ pocket where the primary mutations were located. Considerable losses of potency were observed against protease variants with I84V and I50V mutations for all three inhibitors. The crystal structures revealed an unexpected conformational change in the flap region of I50V protease bound to the analog with the largest P1’ moiety, indicating interdependency between the S1’ subsite and the flap region. Collective analysis of protease-inhibitor van der Waals (vdW) interactions in the crystal structures using principle component analysis indicated I84V mutation underlying the largest variation in the vdW contacts. Interestingly, the principle components were able to distinguish inhibitor identity and relative potency solely based on vdW interactions of active site residues in the crystal structures. Our results reveal the interplay between inhibitor P1’ moiety and primary S1’ mutations, as well as suggesting a novel method for distinguishing the interdependence of resistance through principle component analyses.

---

Human Immunodeficiency Virus (HIV) infects roughly 37 million people globally, with over 2 million new infections and over 1 million AIDS-related deaths occurring each year^1^. After 30 years of research, advances in antiretroviral therapies (ARTs), combinations of small molecule inhibitors, greatly extended life expectancy for those who receive treatment^2,3^. ARTs inhibit critical proteins necessary for viral replication and maturation, most commonly targeting HIV-1 reverse transcriptase and protease. Although ARTs are highly effective in most patients, the current therapies are still an imperfect solution to a complex problem, as HIV-1 can evolve to confer drug resistance through accumulation of mutations^4^. Primary resistance mutations, typically occurring proximal to the active site where the inhibitor binds, are selected early under selective pressure of inhibition and directly affect inhibitor binding and allow the accumulation of additional mutations. Secondary mutations can occur distal to the active site but still indirectly affect substrate processing or inhibitor binding^5,6^. No HIV-1 inhibitor is resistance-proof, and modifying an inhibitor to better tolerate primary resistance mutations may help prevent the accumulation of additional mutations.

HIV-1 protease, a 99-amino acid, homodimeric, aspartyl protease^7,8^ [**Figure 1A**], is essential for viral replication and maturation, making this enzyme an ideal drug target^9,10^. The protease processes twelve unique cleavage sites on viral Gag and Gag-Pol polyproteins to release individual viral proteins required for viral replication and maturation. While the cleavage sites share low amino acid sequence identity, when bound to HIV-1 protease they occupy a consensus volume termed the substrate envelope^11^. We have previously shown that inhibitors that fit within this volume are less prone to resistance as the protease cannot mutate to reduce inhibitor binding without compromising affinity for natural substrates^12^.

**Figure 1.**
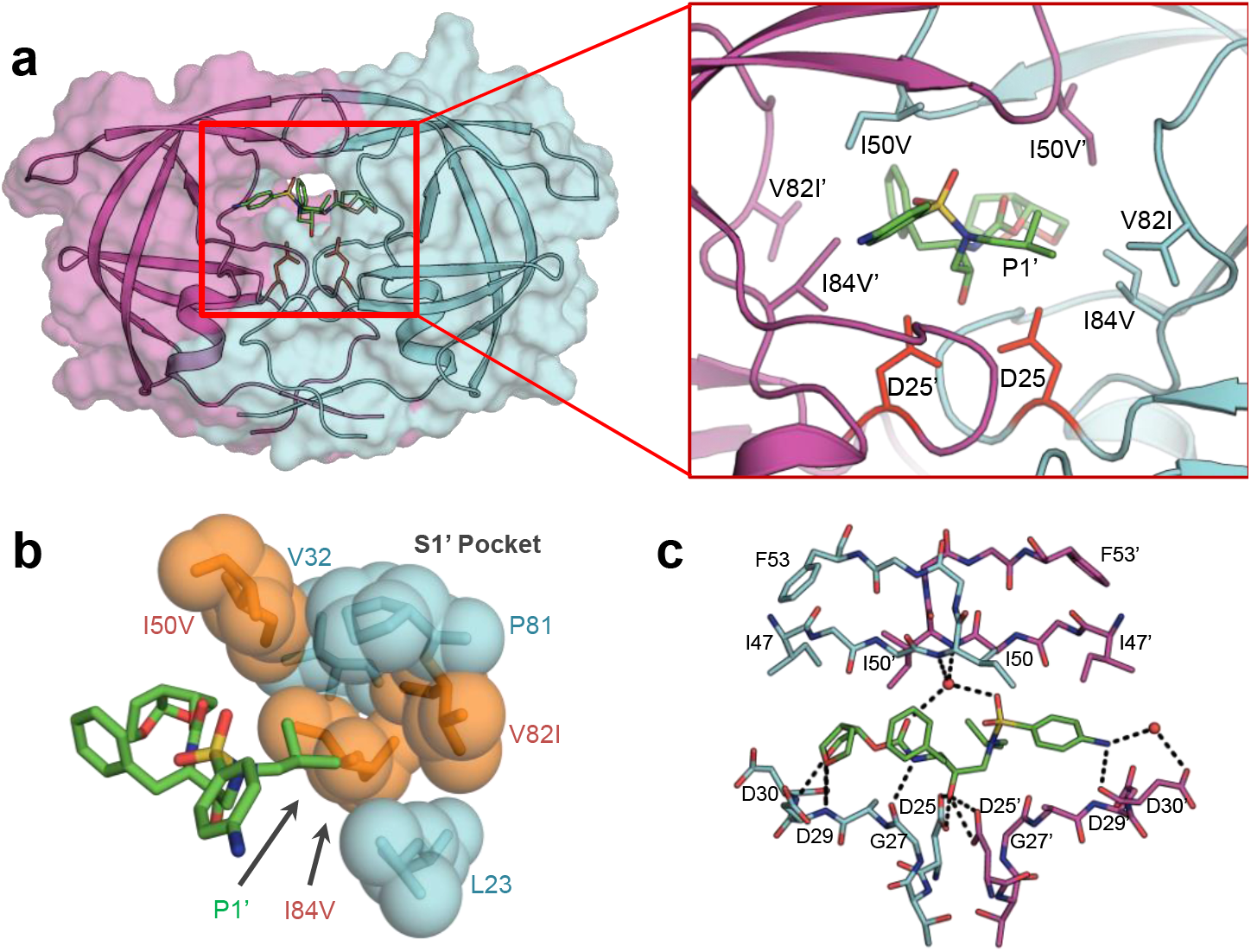
**a)** Co-crystal structure of DRV (green sticks) bound to WT HIV-1 protease. Chain A (cyan) and chain B (magenta) are shown as a cartoon with a transparent surface. D25/D25’ catalytic aspartates (red) are displayed as sticks. **A-Insert)** Residues that contribute to primary drug resistance, shown as sticks. **b)** Spherical representation of residues that make up the S1’ subsite / P1’ pocket. Variable residues are shown in orange. **c)** Hydrogen bonds (black dashes) between DRV and WT HIV-1 protease. Co-ordinated waters are shown as red spheres.

The most potent FDA approved protease inhibitor, darunavir (DRV) [**Figure 2A**], fits well inside the substrate envelope, yet resistance still occurs due to accumulation of mutations in HIV-1 protease in patient isolates^13,14^. DRV is a peptidomimetic inhibitor with four major chemical moieties, denoted P2, P1, P1’ and P2’ [**Figure 2A**]. DRV and analogs have been and continue to be the subject of chemical, viral, structural and dynamics studies^15^. Modifications to the bis-THF P2 moiety of DRV, including fluorine decorations, expansion to tris-THF and fusion into a tri-cyclic group^16-18^ and additional modifications of the P1 and P2’ moiety have been shown to improve potency and resistance profiles^19-21^. Highly mutated clinical isolates with 18 or more mutations in the protease have been identified, and DRV binding to these resistant proteases studied enzymatically and structurally^14,18,22-25^. Most of the patient-derived sequences bear a constellation of secondary mutations as well as primary mutations at the active site that confer DRV resistance, notably including I50V and I84V^26-29^. Mutations at I50 are often selected together with A71V mutation, which is distal from the active site but compensates for the loss of enzymatic fitness^30^. Thermodynamics and structural studies of DRV binding to I50V/L and A71V mutations have revealed significant loss of van der Waals (vdW) contacts between the inhibitor and protease underlying loss of binding affinity^31^.

**Figure 2.**
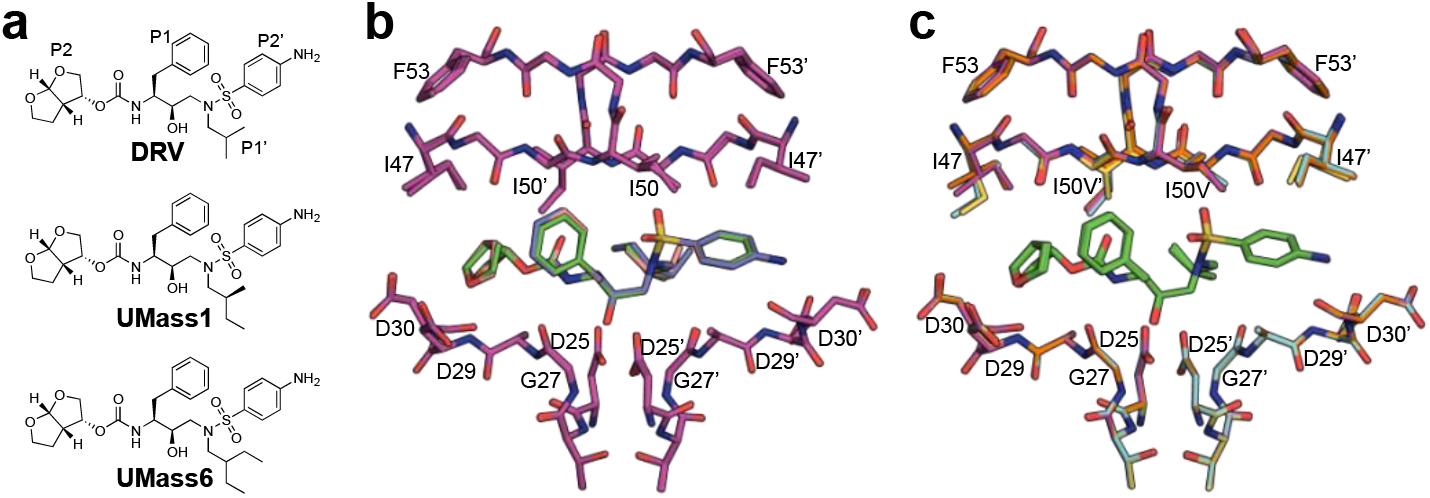
**a)** 2D chemical structure of DRV (with peptidomimetic moieties labeled) and the two DRV analogs (UMass1 and UMass6) with modifications at the P1’ moiety. **b)** WT protease (magenta) inhibited by DRV (green), UMass1 (pink), and UMass6 (purple) **c)** DRV (green) bound to WT (magenta), I84V (cyan), V82I (orange), and I50V (gold) protease variants.

Our previous computational studies based on crystal structures indicated that the P1 and P2 moieties of DRV interacted with HIV-1 protease optimally, even when tested against drug resistance mutations, while the P1’ and P2’ moieties sampled more conformational space during the simulation indicating they could be further optimized^32^. Accordingly, modifying the P1’ moiety to generate DRV analogs with larger hydrophobic groups [**Figure 2A**] leveraged unexploited space in the substrate envelope, improving enzyme inhibition and viral potency against different HIV-1 WT clades and a select panel of drug resistant variants^33^. Crystal structures of these analogs with WT HIV-1 protease, analyzed in comparison to the parent compound DRV, revealed interdependent structural coupling between subsites and suggested modifications at a given inhibitor moiety may affect inhibitor interactions at other locations^34^. The underlying mechanism of how P1’ modifications may affect inhibitor response to resistance mutations surrounding the moiety had not been investigated.

In this work, we investigated the interdependency of potency loss for HIV-1 protease inhibitors that differ at the P1’ moiety (DRV, UMass1, and UMass6) [**Figure 2A**], with WT and 3 variants of HIV-1 protease bearing mutations at residues in the S1/S1’ pocket. The hydrophobic residues I84, V82 and I50 that form the S1’ pocket where the P1’ moiety binds [**Figure 1**] (as well as the S1 pocket, as the protease is a homodimer) are all highly variable, and mutations are implicated in many instances of multidrug resistance, commonly mutating to I84V, V82I, and I50V^13,35,36^. Enzyme inhibition assays were performed and a complete set of 12 crystal structures were determined for the inhibitor-protease combinations. Structural and computational analysis indicated that primary resistance mutations can diminish the potency gain expected from P1’ modifications, by triggering surprisingly large protease rearrangements.

## Results

To elucidate the mechanisms of resistance and potency loss against primary mutations, enzymatic assays and crystal structural analysis were performed with WT HIV-1 protease and 3 variants. All three protease variants had a single mutation that altered the shape of the hydrophobic S1/S1’ pocket of HIV-1 protease: I50V, V82I and I84V. Three inhibitors, DRV with an isobutyl P1’ moiety and 2 analogs in which the P1’ moiety was extended to better fill the substrate envelope, UMass1 with a methylbutyl and UMass6 with an ethlybutyl moiety,^33^ were analyzed.

### Enzymatic Activity of Protease Variants

The enzymatic activity of WT HIV-1 protease and chosen variants (I84V, V82I and I50V) were tested using a natural substrate sequence (MA/CA)^37^ [**Table S1**]. WT NL4-3 HIV-1 protease had a Michaelis-Menten constant, *K*_m_, of 71 ± 7 μM for cleaving this substrate, which was similar to that of I84V and V82I variants (66 ± 4 and 62 ± 4 μM, respectively). The primary resistance mutation I50V is known to reduce catalytic activity,^31^ and while I50V is still catalytically active, the *K*_m_ value is beyond the limit of detection for this assay. A71V, a compensatory mutation that is far from the active site and almost always observed with I50V, restores the functionality to WT level (*K*_m_ = 73 ± 9 μM), as previously reported.^31^ A time-course gel shift assay confirmed the catalytic activity of I50V single mutant, and the rescued activity of I50V/A71V variant in cleaving purified Gag polyprotein [**Figure S1**]. The sustained catalytic activity of proteases across all variants indicates these primary mutations can indeed appear early in drug resistance pathways without compromising substrate cleavage.

### Effect of P1’ Modifications on Inhibition of Variants with Active Site Mutations

The enzyme inhibition constant (*K*_i_) was measured to determine the potency of inhibitors against each HIV-1 protease variant [**Table 1**] using an optimized assay^38^. Although optimized, the lower limit of detection is about 5 pM and all 3 inhibitors were too potent to obtain a reliable value against WT protease. The V82I mutation did not confer any measurable resistance for the inhibitors, with the *K*_i_ staying below 5 pM. The I84V mutation caused a reduction in potency, with the *K*_i_ increasing to around 25 pM for DRV and UMass1, and approximately half of that for UMass6. Thus, the inhibitor with the largest P1’ moiety performed 2-fold better than the other inhibitors against I84V variant. The I50V mutation was more deleterious with *K*_i_ values increasing two orders of magnitude relative to WT protease. Interestingly, UMass6 performed slightly worse than DRV and UMass1 (*K*_i_ = 146 ±11 versus 117 ± 6 and 131 ± 8 pM, respectively). Although both I84V and I50V mutations cause the same reduction in side chain size, these two mutations had the opposite effect on the change in potency of UMass6 compared to DRV and UMass1.

**Table 1.** Enzyme inhibition constant, *K*_i_ (pM), of inhibitors against WT protease and variants with primary resistance mutations, measured using an optimized enzymatic assay.

| Inhibitor | WT | I84V | V82I | I50V | I50V/A71V |
| --- | --- | --- | --- | --- | --- |
| DRV | < 5 | $25.6 \pm 5.6$ | < 5 | $117.2 \pm 5.8$ | $74.5 \pm 5.6$ |
| UMass1 | < 5 | $26.1 \pm 3.7$ | < 5 | $131.3 \pm 8.2$ | $110.3 \pm 8.8$ |
| UMass6 | < 5 | $12.8 \pm 3.1$ | < 5 | $146.2 \pm 10.7$ | $100.0 \pm 9.9$ |

### Structural Rearrangements Underlying DRV Susceptibility to Primary Resistance Mutations

To determine the structural basis of observed potency changes due to primary resistance mutations around the P1’ moiety, each protease variant was co-crystalized with each inhibitor resulting in a set of 12 crystal structures [**Table S2**]. All structures were solved in the same space group with a resolution of 2 Å or better, affording detailed investigation of atomic interactions. We determined the crystal structures of DRV, UMass1 and UMass6 bound to WT HIV-1 protease of NL4-3 strain, which was also used in the enzymatic assays above.

When DRV and other peptidomimetic inhibitors bind to HIV-1 protease, the hydroxyl moiety is centered between the catalytic aspartates (D25/D25’) and the inhibitor makes a number of additional hydrogen bonding interactions with the backbone nitrogen and oxygen atoms of residues that line the active site [**Figure 1C**]. DRV also makes a key water-mediated interaction with the backbone nitrogen of residue 50, located at the tip of each flap. DRV’s P1 and P1’ moieties are hydrophobic and interact with hydrophobic residues/pockets in the hinge of the flaps. Inhibitor potency diminishes if any of these polar or non-polar interactions are perturbed.

Comparison of DRV-bound crystal structures revealed the structural rearrangements in response to single mutations at the active site. In agreement with the maintained potency against V82I variant, the hydrogen bonding and packing of DRV at the active site was conserved in the V82I crystal structure [**Figure 2B and Table 2**]. The additional steric bulk of the side chain due to V82I is not directed towards the inhibitor but instead is solvent exposed, which does not perturb inhibitor binding. As in the V82I variant, the binding mode and hydrogen bonds of DRV were not altered in I84V and I50V structures [**Figure 2B**]. However, van der Waals (vdW) interactions with residues 84 and 84’ decreased due to I84V mutation, which was in part compensated by increased packing against I47 and V82’ [**Table 2**]. In the I84V variant, residues V32’ and L23’ in chain B underwent a side chain conformer change relative to WT protease [**Figure S2].** In addition, residue I47 was found in an alternative rotamer in chain A, causing increased inhibitor interactions with that residue. This asymmetric compensation indicates subtle alterations in the overall packing of the inhibitor at the active site, which reduced the effect of I84V mutation on potency. A similar repacking was also observed in the I50V–DRV structure, where increased packing against I47 in both chains compensated lost vdW interactions at residue 50. Our previous computational analysis had indicated I47 as a major modulator of DRV-protease interactions^39^. Thus, the structures explain why the V82I single mutation did not confer resistance to DRV, and in agreement with previous studies reveal the rearrangements in vdW packing around DRV underlying susceptibility to I50V^31,40^.

**Table 2.** Change in intermolecular van der Waals (vdW) interactions between inhibitor and protease active site residues relative to WT complex. Only residues with changes greater than 0.40 kcal/mol are listed. Primed residue numbers correspond to chain B. Decrease and increase in per residue contacts with respect to WT protease are colored blue to red, respectively.

| Resi # | I84V_DRV | I84V_UMass1 | I84V_UMass6 | V82I_DRV | V82I_UMass1 | V82I_UMass6 | I50V_DRV | I50V_UMass1 | I50V_UMass6 |
| --- | --- | --- | --- | --- | --- | --- | --- | --- | --- |
| 30 | 0.28 | -0.14 | -0.83 | 0.46 | 0.05 | -0.44 | 0.00 | -0.01 | -0.48 |
| 47 | 1.22 | 1.07 | 1.12 | -0.02 | -0.04 | 1.12 | 1.19 | -0.08 | -0.18 |
| 48 | 0.49 | 0.10 | 0.22 | 0.14 | 0.13 | 0.26 | 0.34 | 0.05 | -0.18 |
| 49 | 0.33 | -0.02 | 0.08 | 0.06 | 0.10 | 0.25 | 0.19 | 0.27 | -0.44 |
| 50 | -0.50 | -0.60 | -0.53 | -0.11 | -0.04 | 0.13 | -1.70 | -1.32 | -1.94 |
| 84 | -1.17 | -1.10 | -1.17 | -0.12 | 0.15 | -0.01 | -0.05 | 0.27 | -0.31 |
| 23' | -0.60 | -0.33 | -0.25 | -0.06 | -0.17 | -0.11 | 0.10 | -0.09 | -0.07 |
| 25' | -0.41 | -0.31 | -0.37 | 0.10 | 0.01 | -0.11 | -0.08 | 0.04 | -0.30 |
| 28' | -0.55 | -0.34 | -0.28 | 0.06 | 0.29 | 0.24 | 0.02 | 0.40 | -0.47 |
| 47' | -0.36 | -0.47 | -0.42 | 0.20 | 0.04 | -0.02 | 0.49 | 0.09 | 0.41 |
| 48' | -0.16 | -0.03 | 0.03 | -0.01 | 0.16 | 0.19 | 0.15 | 0.15 | 0.44 |
| 49' | -0.09 | -0.07 | 0.07 | -0.03 | 0.09 | 0.19 | 0.18 | 0.16 | 0.61 |
| 50' | -0.08 | 0.16 | -0.03 | -0.09 | 0.11 | 0.03 | -0.84 | -0.44 | 0.17 |
| 82' | 1.17 | 1.03 | 0.83 | 0.23 | 0.41 | 0.27 | 0.10 | 0.15 | 0.21 |
| 84' | -1.82 | -1.61 | -1.54 | -0.03 | -0.03 | -0.02 | 0.03 | -0.23 | -0.21 |

### Structural Response to Modifying P1’ Moiety Against Primary Resistance Mutations

The larger P1’ moieties of UMass1 and UMass6 were designed to improve hydrophobic packing and increase vdW contacts with the protease while still staying within the substrate envelope^33^. Accordingly, the total vdW intermolecular interactions between the inhibitor and protease increased by 1.6 and 4.7 kcal/mol, respectively for UMass1 and UMass6, as the P1’ moiety increased in size [**Table S3**]. In the V82I structure, while DRV did not gain significant interactions compared to WT complex, UMass1 and UMass6 gained approximately 2 kcal/mol in vdW energy as their P1’ moieties directly interact, but not clash, with the sterically larger side chain. Thus, although the *K*_i_ values are below the limit of detection, structural analysis suggests that V82I mutation might confer hyper-susceptibility to UMass1 and UMass6 inhibitors; thus unlike with DRV, the V82I mutation is not likely to be selected under the pressure of these inhibitors.

The co-crystal structures of UMass1 bound to WT protease and variants were very similar to those of DRV [**Figure 2C**], albeit with subtle alterations. Against I84V and I50V mutations, which are located directly at the pocket where P1’ moiety binds, UMass1 experienced a similar reduction in potency as DRV [**Table 1**]. The UMass1–I84V structure displayed the same phenomenon of asymmetric inhibitor repacking as with DRV [**Table 2**]. The additional methyl group in UMass1 is oriented away from the steric space provided by the I84V mutation but still makes additional contacts with P81 and V82. When UMass1 bound I50V, there was a reduction in vdW interactions at residue 50 and 50’ as seen for DRV, but no compensation at residue 47. Instead, repacking of UMass1 increased against G49 and I84 in chain A, and residues 28–30 in chain B. However, this alternate compensation was more distributed and subtle, which may underlie the slightly worse *K*_i_ of UMass1 compared to DRV against I50V protease. Thus, the crystal structures indicate that although the potency loss against I50V is similar for DRV and UMass1, the underlying structural changes and repacking at the active site are distinct.

### I50V Mutation Induces Conformational Changes in Both UMass6 and Protease Flaps

UMass6 has an even bulkier P1’ moiety and binds WT protease similar to DRV and UMass1 [**Figure 2C**] but with enhanced overall vdW interactions (−87.8 kcal/mol compared to −83.1 and −84.7, respectively; **Table S3**). Against I84V mutation, enhanced packing at the P1’ moiety of UMass6 resulted in overall better interactions and potency. In UMass6, the methyl groups on both branches of the isobutyl P1’ moiety help UMass6 better accommodate the steric space provided by the I84V mutation, leading to a 2-fold improvement in *K*_i_ compared to the other two inhibitors.

Contrary to the case of I84V mutation, UMass6 bound to the I50V variant with the lowest potency. As with UMass1, extending the P1’ moiety further resulted in a worse potency against I50V mutation than DRV. Unexpectedly, the P1’ moiety of UMass6 in I50V protease structure adopted a completely different conformation compared to the other 11 structures [**Figure 3A**]. In addition to this inhibitor’s unique binding conformation, the flaps in I50V protease underwent rather large-scale changes: In chain A, I47 assumed a rare rotamer to maintain interaction with the sterically smaller I50V [**Figure S2**]. Interestingly, in chain B, residue 50’ underwent a shift away from the inhibitor as well as a T80 residue flip. Residue T80 is one of 27 invariant residues and is necessary for flap flexibility^41^. Conformational changes at residues P79-T80-P81 have been tied to flap-tip curling in both MD simulations and NMR structures^42^. The mutated residues 50 and 50’ are located at the tips of the protease flaps. When the protease flaps close around an inhibitor, the backbone polar atoms of each residue 50 form a hydrogen bonding interaction. Because DRV and natural protease substrates are asymmetric and HIV-1 protease is actually a pseudo-homodimer, residue 50 interactions at the flaps are commonly found in the same specific asymmetric conformation. Surprisingly, when UMass6 was bound to I50V, the flaps underwent a conformational change relative to the typically observed structures [**Figure 3B-C**]. This flip caused the flaps to bow outward [**Figure 3C**] away from UMass6, further reducing vdW contacts. Interestingly, the resulting vdW losses of UMass6 with residue 50 due to the I50V mutation were completely asymmetric, with a loss similar to DRV and UMass1 in chain A but almost no loss in chain B [**Table 2**]. However, there was no compensation at residue 47 interactions in either chain, due to the different conformer of I47, which resulted in a greater overall loss in vdW interactions, in agreement with the greatest potency loss observed for UMass6 against I50V mutation.

**Figure 3.**
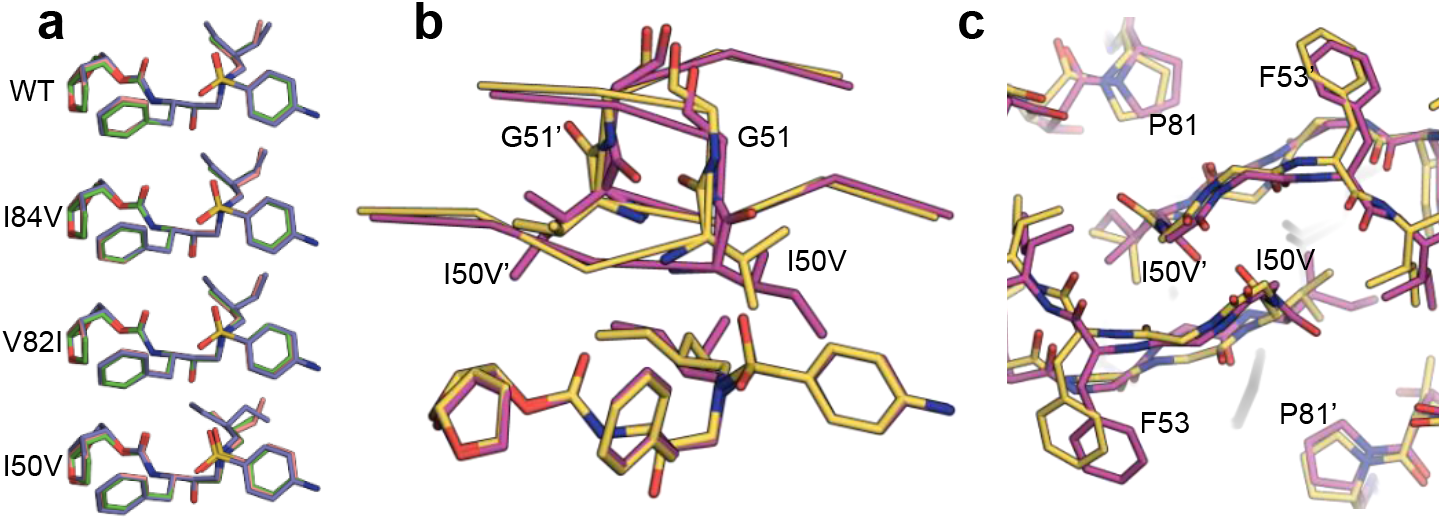
**a)** Alignment of inhibitors in all 12 crystal structures, grouped by protease variant. DRV (green), UMass1 (pink) and UMass6 (purple) in WT, I84V, V82I, and I50V protease crystal structures. The P1’ moiety of UMass6 (purple) bound to the I50V variant exists in an alternate conformation. **b)** Backbone interaction of residues at the flap tips in WT-UMass6 (magenta) and I50V-UMass6 (gold) structures. Residues 50 and 51 are shown as sticks and residues 47-53 are shown as ribbons. **C)** Top view of the flap region of WT-UMass6 (magenta), and I50V-UMass6 (gold), shown as sticks.

Although the position of most of the flap residues were altered in the UMass6–I50V structure relative to WT complex, the typical hydrogen bond distances were not significantly perturbed [**Figure S3, Table S4**]. However, the flipping of the flaps caused a noticeable shift in the location of the so-called flap water [**Figure S3**], which in turn caused the sulfonyl group to shift toward the flaps to maintain H-bond distances. The shift of the sulfonyl group, which is fused to the aniline P2’ moiety, weakened the bond formed by the conjugated water interacting with the side chain of D30 by making the overall distance longer.

The structural rearrangements of the protease in the I50V–UMass6 structure were quantified and visualized by distance-difference matrices [**Figure S4**]. Distance-difference matrices measure the internal distances between alpha carbon atoms of every residue pair in a structure, then compare each distance to that in a reference structure, which in our case is the inhibitor bound to WT protease. This method detects conformational changes due to a mutation without any structural superposition bias. Comparison of the variant structures to their corresponding WT structures shows the discrepancy between the I50V–UMass6 and all the other 11 structures [**Figure S4].** Mapping the distance-differences onto the 3D structure revealed that the structural changes due to I50V in the UMass6-bound protease propagated throughout the protein structure [**Figure 4**].

**Figure 4.**
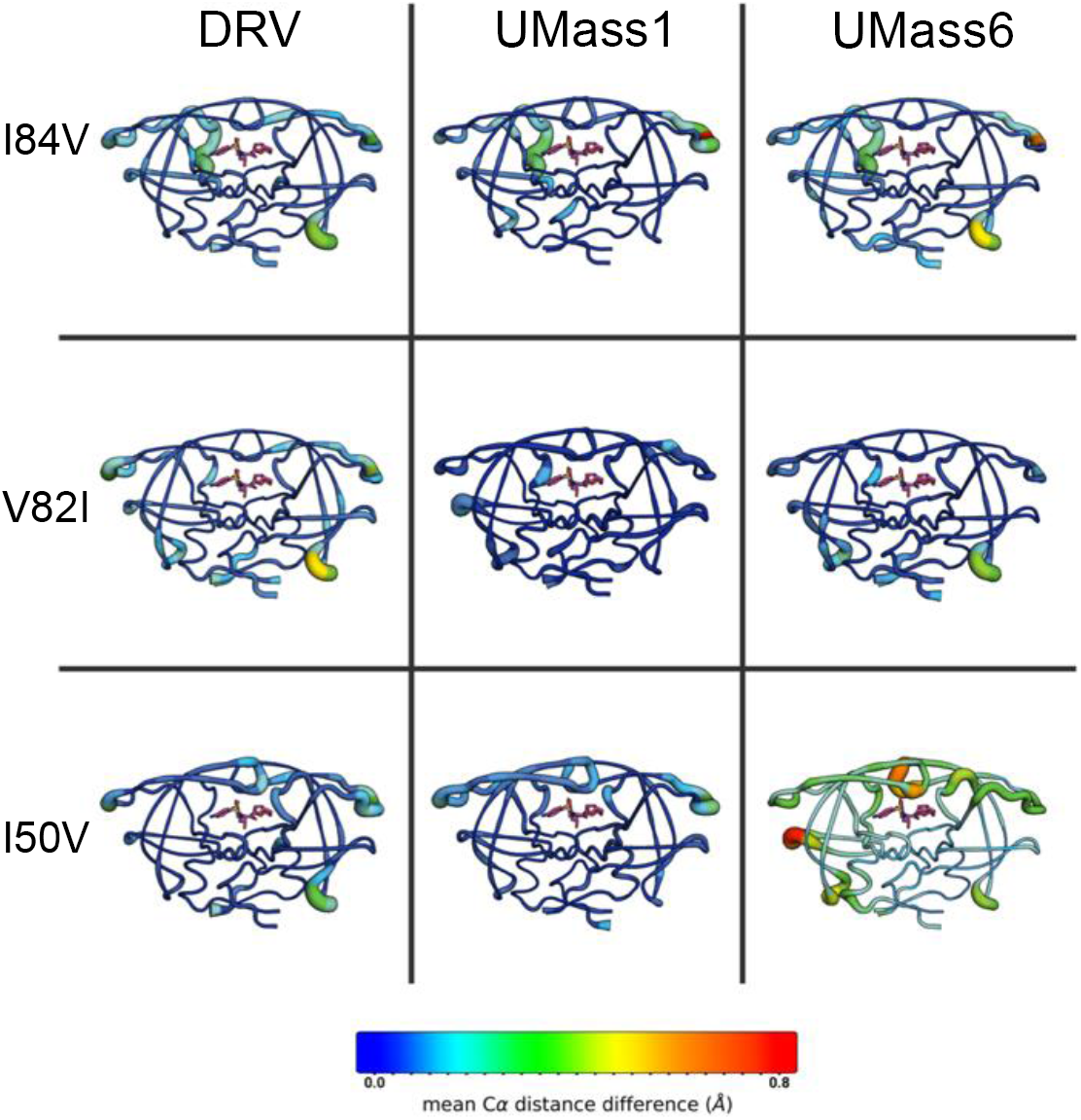
Cartoon-putty diagrams depicting the mean changes in C-alpha distance differences relative to WT protease crystal structure. Tube thickness and warm colors indicate larger differences.

### Principle Component Analysis (PCA) of Residue-Specific Interactions

The inhibitor–protease vdW interactions in the 12 crystal structures were subjected to principal component analysis (PCA) to investigate the variance therein due to primary resistance mutations and inhibitor P1’ moiety modifications. The 198×198 covariance matrix was calculated based on the residue-wise vdW interactions. The first three principal components (PCs) accounted for 83% of the observed variance, with the first PC (PC1) accounting for approximately half (43%) of the variance [**Figure S5A].** Location and clustering of structures along the PCs reveal what causes the variations in inhibitor–protease interactions among these structures.

The PC1 separated the structures mainly according to the protease variant, rather than inhibitor identity [**Figure 5].** Significantly, structures containing the I84V primary resistance mutation of all 3 inhibitors clustered away from the remaining structures. Analysis of per residue contributions to the ordering along the principal component indicated that the spread along the PC1 was largely determined by changes in vdW interactions between the inhibitor and residue 84. This is in agreement with our previous work that showed that mutations at residue 84 significantly alter the pattern of vdW contacts in a panel of drug resistant variants.^43^ Additionally, contacts between I47 and the inhibitor contributed significantly to the spread along PC1, which can assume a different side chain conformer and compensate for loss of interactions at residue 84 as explained above.

**Figure 5.**
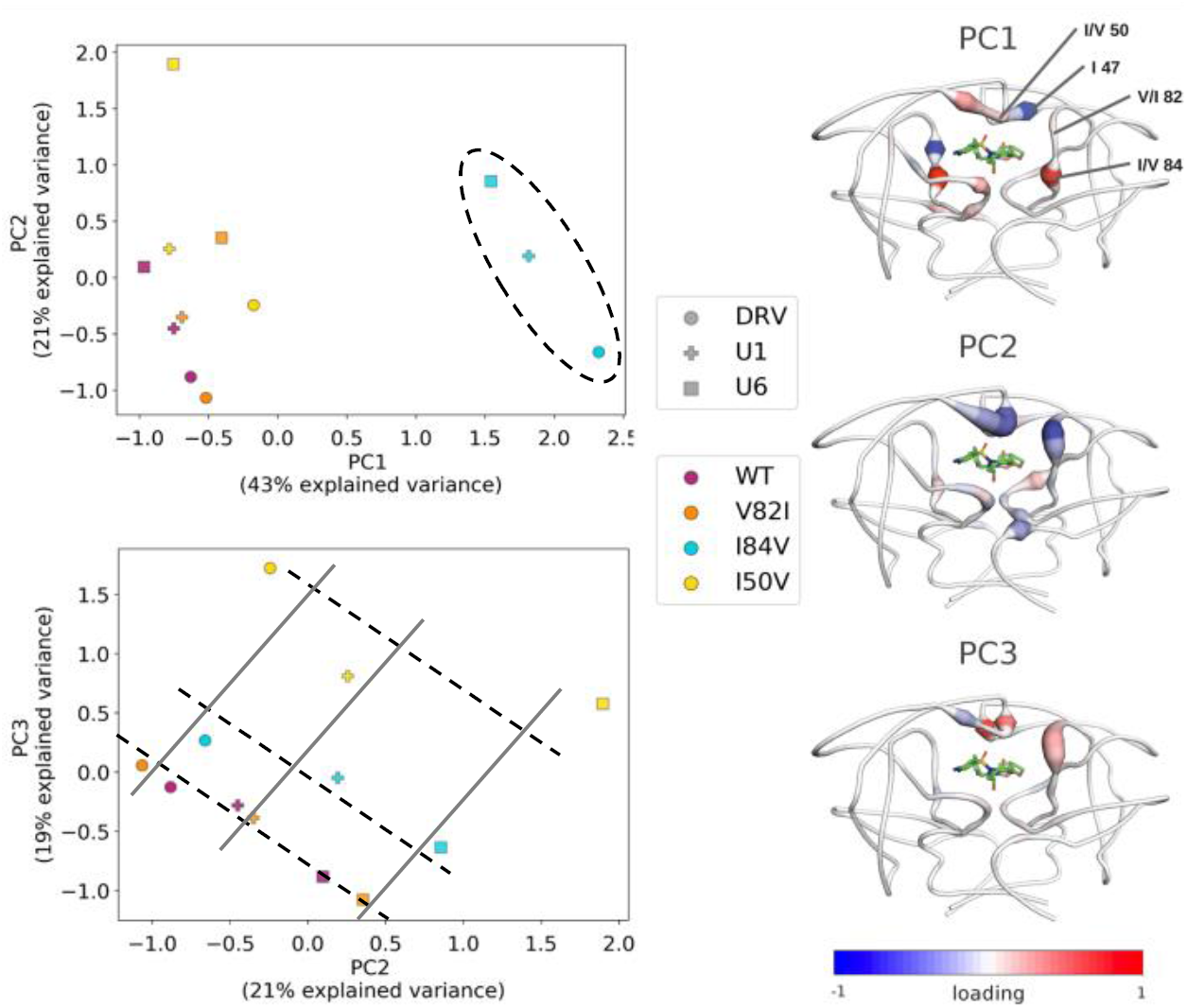
Principle component analysis of vdW interactions in the 12 protease-inhibitor complexes. The protease-inhibitor pairs plotted according to first and second principle components, PC1 versus PC2 (upper panel) and PC2 versus PC3 (lower panel). The dashed oval and parallel lines are shown only to guide the eye. The contribution of individual residues to the PCs according to the loading vector are depicted on protease backbone structure using cartoon-putty diagrams, where tube thickness indicates higher weights and warm colors are positive loading and cold colors are negative loading. Residues with the highest weights are labeled on the cartoon-putty for PC1.

The second and third principal components (PC2 and PC3) together accounted for 40% of the observed variance in inhibitor–protease vdW interactions. PC2 separated structures mostly according to inhibitor identity, rather than protease variant [**Figure 5**]. Inspection of the loading of PCs showed that variance at residues 84 and 50’ contributed significantly to the ordering of structures along PC2, indicating modification of inhibitor P1’ moiety caused alteration in packing against these residues [**Figure S5B**]. The contribution of residue 50 in chain A had the same direction for both PC2 and PC3 whereas the contributions of 84 and 50’ had the opposite direction. Contacts between the inhibitor and residues 84 and 50’ changed significantly in the presence of a small P1’ moiety but were less effected when the P1’ moiety was larger. Overall, the PCA captured the asymmetric compensation of vdW interactions observed in the crystal structures, and quantified the contribution of residue-specific interactions to the overall variance in the dataset.

Strikingly, plotting PC2 and PC3 against each other enabled separating the 12 crystal structures both according to inhibitor identity and potency [**Figure 5**]. All 6 inhibitor-protease pairs with <5 pM affinity clustered on a line defined by a linear combination of PC2 and PC3. Intriguingly, the I84V and I50V complexes fell on separate but parallel lines, together defining a set of 3 roughly parallel lines on the PC2-PC3 plane. These 3 lines were ordered according to experimentally measured potency, with the I84V complexes nearer to the high affinity line, followed by the I50V complexes. Additionally, 3 other lines perpendicular to these 3 parallel lines separated the inhibitor-protease pairs according to inhibitor identity. The inhibitors were also ordered, according to the chemical similarity of their P1’ moieties as DRV-UMass1-UMass6. Thus, the collective analysis of the set of 12 crystal structures using PCA enabled clustering inhibitor identity and potency based solely on per residue vdW contacts.

## Discussion

We have investigated here the interplay between modification of the P1’ moiety in the DRV scaffold, and primary protease mutations at residues forming the active site pocket where this moiety binds. To improve DRV potency and resistance profile, the P1’ moiety was modified to optimally fill the substrate envelope and increase contacts with hydrophobic residues in this pocket. These larger, flexible P1’ groups were previously shown to increase potency in cellular assays, specifically against a panel of WT clades and drug resistant variants^33^. Our results here validated that larger P1’ moieties were effective against I84V and V82I mutations, but an unexpected alternative mechanism of resistance was uncovered against the I50V mutation. The basic idea that a larger P1’ moiety would help the inhibitor retain better interactions upon shortening of a side chain is not necessarily correct. Rather than a simple alteration in interactions at this pocket, there was an overall and asymmetric rearrangement of the vdW interactions throughout the active site pocket due to either I84V or I50V mutation. As intended, the bulkier P1’ moiety of UMass6 helped retain better potency against I84V, but the reverse was the case for I50V mutation. Thus, although both I84V and I50V are primary resistance mutations with the same side chain change in the S1’ pocket, the interplay between P1’ moiety and rearrangements of inhibitor interactions were distinct, and moreover contrasting for these two mutations.

The I50V mutation is commonly observed in multi-mutant variants that are resistant to DRV, and has previously been investigated^23,29^. This mutation disrupts the inter-protease hydrophobic packing involving the flap tips and residues V32, I47, and I84, which together stabilize the closed conformation of protease. When residue 50 is truncated to a valine, the flaps most likely cannot efficiently close around the bound ligand, either inhibitor or substrate, which results in reduction in catalytic activity as well as loss in inhibition. The compensatory mutation in I50V/A71V double mutant restores both catalytic activity and increases inhibitor potency for all 3 inhibitors, in agreement with the idea that flap closing is key for both substrate processing and inhibition. Unlike DRV and UMass1, the P1’ group of UMass6 is large enough to be able to contribute a methyl group to interact with the destabilized hydrophobic interactions involving the flap tips. However, this interaction affects the orientation of V50, which to avoid an unfavorable side chain rotamer, induces flipping of the flaps and a rather unforeseen conformational rearrangement. A similar but distinct flap rearrangement was previously observed in response to a coevolution mutation in Gag substrate in the I50V/A71V variant, which enhances vdW interactions with the substrate^44^. Hence the relative flexibility of flaps in this variant might be able to enhance interactions with substrates while weakening inhibitor binding, thus contributing to conferring resistance.

Unlike the other two mutations, V82I did not confer measurable resistance against the inhibitors tested. However, crystal structures suggested that the intermolecular interactions are improved for UMass1 and UMass6 relative to WT protease, unlike for DRV. Thus, the bulkier P1’ groups in UMass inhibitors might cause hyper-susceptibility to V82I and prevent early selection of V82I as a primary resistance mutation to result in a different resistance pathway compared to DRV. Although a larger P1’ moiety increases favorable nonpolar interactions with HIV-1 protease, groups larger than that in UMass1 may act as a selective pressure forcing I50V to become a dominant variant, especially at early times during the accumulation of resistance mutations under the selective pressure of inhibitors. Larger P1’ moieties may induce I50V selection as the first step in a pathway of resistance. Thus, despite high similarity, inhibitors with P1’ modifications may select distinct mutational patterns of resistance.

The set of 12 high-resolution crystal structures comprising 3 analogous inhibitors bound to WT protease and 3 variants allowed a collective analysis of inhibitor-protease interactions in modulating potency and resistance. PCA not only identified the key residues that determined the variance in inhibitor packing but also captured the asymmetry in this variance. Interestingly, the major principle components were able to categorize the inhibitor-protease pairs according to inhibitor type and potency. This suggests residue-specific vdW interactions may serve as “fingerprints” to categorize and classify inhibitors bound to protease. Moreover, certain linear combinations of principal components might be indicators or even predictors of inhibitor potency. Regardless, our comprehensive analysis reveals the interplay between inhibitor modifications and structural response of the target with primary mutations underlying resistance.

## Materials and Methods

### Protease gene construction

Protease gene construction was carried out as previously described^44,45^. The NL4-3 strain has four naturally occurring polymorphisms in the protease relative to the SF-2 strain^15,46^. In short, the protease variant genes (I50V, V82I, I84V) were constructed using QuikChange site-directed mutagenesis (Genewiz) onto NL4-3 WT protease on a pET11a plasmid with codon optimization for protein expression in Escherichia coli. A Q7K mutation was included to prevent autoproteolysis^47^.

### Protein expression and purification

The expression, isolation, and purification of WT and mutant HIV-1 proteases used for the kinetic assays and crystallization were carried out as previously described^44,45^. Briefly, the gene encoding the HIV protease was subcloned into the heat-inducible pXC35 expression vector (ATCC) and transformed into E. coli TAP-106 cells. Cells grown in 6L of Terrific Broth were lysed with a cell disruptor and the protein was purified from inclusion bodies^48^. The inclusion body centrifugation pellet was dissolved in 50% acetic acid followed by another round of centrifugation to remove impurities. Size exclusion chromatography was used to separate high molecular weight proteins from the desired protease. This was carried out on a 2.1-L Sephadex G-75 superfine (Sigma Chemical) column equilibrated with 50% acetic acid. The cleanest fractions of HIV protease were refolded into a 10-fold dilution of 0.05 M sodium acetate at pH 5.5, 5% ethylene glycol, 10% glycerol, and 5 mM DTT. Folded protein was concentrated down to 0.5-3mg/mL and stored. This stored protease was used in K_M_ and K_i_ assays. For crystallography, a final purification was performed with a Pharmacia Superdex 75 FPLC column equilibrated with 0.05 M sodium acetate at pH 5.5, 5% ethylene glycol, 10% glycerol, and 5 mM DTT. Protease fractions purified from the size exclusion column was concentrated to 1-2 mg/mL using an Amicon Ultra-15 10-kDa device (Millipore) for crystallization.

### Enzymatic Assays to Determine K_*m*_ and K_*i*_

The *K*_*m*_ and *K*_*i*_ Assays were carried out as previously described^37,38^. In the *K*_*m*_ assay, a 10-amino acid substrate containing the natural MA/CA cut site with an EDANS/DABCYL FRET pair was dissolved in 8% DMSO at 40nM and 6% DMSO at 30 nM. The 30 nM of substrate was 4/5^th^ serially diluted from 30 nM to 6 nM, including a 0 nM control. HIV protease was diluted to 120 nM and, using a PerkinElmer Envision plate reader, 5 μL was added to the 96-well plate to obtain a final concentration of 10 nM. The fluorescence was observed with an excitation at 340 nm and emission at 492 nm and monitored for 200 counts, for approximately 23 minutes. FRET inner filter effect correction was applied as previously described^49^. Corrected data was analyzed with Prism7.

To determine the *K*_*i*_, in a 96-well plate, each inhibitor was 2/3 serially diluted from 3000 pM to 52 pM, including a 0 pM control, and incubated with 0.35 nM protein for 1 hour. A 10-amino acid substrate containing an optimized protease cut site with an EDANS/DABCYL FRET pair was dissolved in 4% DMSO at 120 μM. Using the Envision plate reader, 5 μL of the 120 μM substrate was added to the 96-well plate to a final concentration of 10 μM. The fluorescence was observed with an excitation at 340 nm and emission at 492 nm and monitored for 200 counts, for approximately 60 minutes. Data was analyzed with Prism7.

### Protein Crystallization

Discovery of the condition producing reproducible co-crystals of DRV with NL4-3 WT protease was achieved using the JCSG+ screen (Molecular Dimensions), in well G11, consisting of 2 M Ammonium Sulfate with 0.1 M Bis-Tris-Methane Buffer at pH 5.5 with a protease concentration of 1.9 mg/mL with 3-fold molar excess of DRV and mixed with the precipitant solution at a 1:2 ratio. After optimization, all subsequent combinations of co-crystals were grown at room temperature by hanging drop vapor diffusion method in a 24-well VDX hanging-drop trays (Hampton Research) with protease concentrations between 1.0 to 2.4 mg/mL with 3-fold molar excess of inhibitors set the crystallization drops with the reservoir solution consisting of 23-24% (w/v) Ammonium sulfate with 0.1 M bis-Tris-methane buffer at pH 5.5 set with 2 μL of well solution and 1 μL protein and microseeded with a cat whisker. Diffraction quality crystals were obtained within 1 week. As data was collected at 100 K, cryogenic conditions contained the precipitant solution supplemented with 25% glycerol.

### Data Collection and Structure Solution

Diffraction data were collected and solved as previously described^44,50^. Diffraction quality crystals were flash frozen under a cryostream when mounting the crystal at our in-house Rigaku_Saturn944 X-ray system. The co-crystal diffraction intensities were indexed, integrated, and scaled using HKL3000^51^. Structures were solved using molecular replacement with PHASER^52^. Model building and refinement were performed using Coot^53^ and Phenix^54^. Ligands were designed in Maestro and the output sdf file was used in the Phenix program eLBOW^55^ to generate the cif file containing atomic positions and constraints necessary for ligand refinment. Iterative rounds of crystallographic refinement were carried out until convergence was achieved. To limit bias throughout the refinement process, five percent of the data were reserved for the free R-value calculation^56^. MolProbity^57^ was applied to evaluate the final structures before deposition in the PDB. Structure analysis, superposition and figure generation was done using PyMOL.^58^ X-ray data collection and crystallographic refinement statistics are presented in the Supporting Information [**Table S2**].

### Structural Analysis and PCA

Distance-difference matrices were generated for each inhibitor-mutant protease pair to reveal structural changes relative to that inhibitor bound to wild-type protease, as previously described^59^. Briefly, distances between all Cα pairs in the mutant structure were calculated as an *N*x*N* matrix (N = 198 residues for HIV-1 protease), and then those corresponding distances in the wild-type structure were subtracted to construct the distance difference matrix. The mean deviation from the WT structure for each residue was then calculated by taking the average of the absolute value of all the *N* distance differences involving that residue, and the backbone structure was represented as a cartoon-putty with increasing thickness and warmer color for increasing deviation.

To calculate the intermolecular van der Waals (vdW) interaction energies the crystal structures were prepared using the Schrodinger protein preparation wizard^60^. Hydrogen atoms were added, protonation states determined and the structures were minimized. Subsequently forcefield parameters were assigned using the OPLS2005 forcefield^61^. Interaction energies between the inhibitor and protease were estimated using a simplified Lennard-Jones potential, as previously described^62^. For each protease residue, the change in vdW interactions relative to WT complex was also calculated for the mutant structures. PCA of the data matrix was performed as described earlier.^43^ For the calculation of the principal components, the implementation of PCA in scikit-learn was used^63^. Briefly, the intermolecular vdW interaction energy with the inhibitor for each residue in a given structure was calculated for each of the 12 structures to yield a 198×12 matrix. The dimensionality of the data set was then reduced using PCA to identify orthogonal linear combinations of variables, or principal components (PCs), that best account for the variance in the data. The PCs are ordered starting from first PC according to the greatest variance represented in the data, and contribution of the original variables to a given PC is represented by the loading vector.

## Acknowledgements

This research was supported by NIH P01 GM109767. We would like to thank G. Nachum for performing the Gag polyprotein cleavage assay, and W. Royer, B. Kelch and B. Hilbert for helpful discussions.

## Table of Contents Graphic

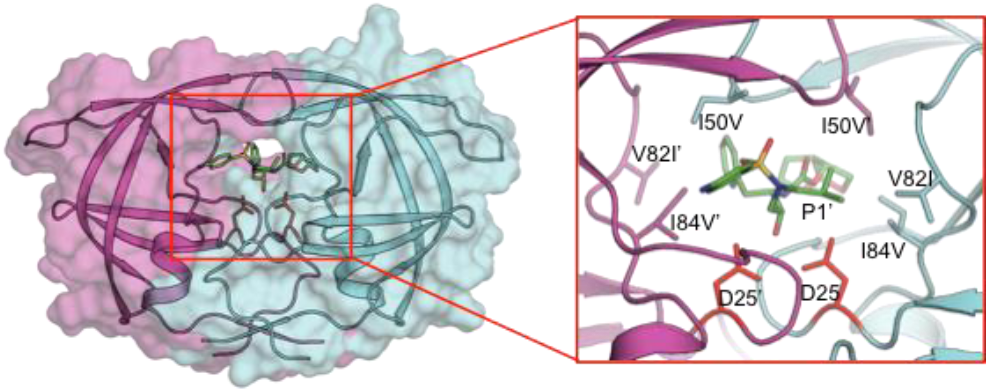

